# Sensorimotor memories influence movement kinematics but not associated tactile processing

**DOI:** 10.1101/2023.03.18.533257

**Authors:** Marie C. Beyvers, Dimitris Voudouris, Katja Fiehler

**Author notes:** Corresponding author at: Department of Experimental Psychology, Justus Liebig University Giessen, Otto-Behaghel-Strasse 10F, 35394 Giessen, Germany. *e-mail address* (K. Fiehler).

## Abstract

When interacting with objects, we often rely on visual information. However, vision is not always the most reliable sense for determining relevant object properties. For example, when the mass distribution of an object cannot be inferred visually, humans may rely on predictions about the object’s dynamics. Such predictions may not only influence motor behavior but also associated somatosensory processing, as sensorimotor predictions lead to reduced tactile sensitivity during movement. We examined whether predictions based on sensorimotor memories influence grasping kinematics and associated tactile processing. Participants lifted an object of unknown mass distribution and reported whether they detected a tactile stimulus on their grasping hand during the lift. In Experiment 1, the mass distribution could change from trial to trial, whereas in Experiment 2, we intermingled longer with shorter parts of constant and variable mass distributions, while also providing implicit or explicit information about the trial structure. In both experiments, participants grasped the object by predictively choosing contact points that would compensate the mass distribution experienced in the previous trial. Tactile suppression during movement, however, was invariant across conditions. These results suggest that predictions based on sensorimotor memories can influence movement kinematics but may not affect associated tactile perception.

**Public significance statement:** To perform a goal-directed movement, such as grasping an object, humans combine the available sensory information with predictions about the prevailing dynamics. Sensorimotor predictions also lead to a decrease of movement-related tactile signals, a phenomenon termed tactile suppression. Tactile suppression is supposed to rely on a dynamic weighting of sensory feedback and predictive signals. When sensory feedback is not reliable, reliance on memory-based predictions may be desired. Here we show that motor behavior is influenced by predictions based on sensorimotor memories, but associated tactile processing appears to be robust.

## Introduction

Successful human-world interactions require processing of relevant sensory signals. For example, when grasping an object, humans choose grasping points based on visual information about the object’s size (Gordon et al., 1991), orientation (Voudouris et al., 2013), shape (Kleinholdermann et al., 2007), or surface material (Klein et al., 2021). However, vision is not always the most reliable sense to work out relevant object properties. Indeed, although object mass can be inferred from visual information, as mass typically increases with object size, there are occasions when vision misleads our judgements, such as the size-weight illusion (Flanagan & Beltzner, 2000). Likewise, although an object’s mass distribution can be inferred by the object’s shape, some objects that appear to be symmetric may actually have an asymmetric mass distribution. When grasping objects with properties that cannot be reliably inferred from vision, such as of unknown mass distribution, humans *appear* to adopt a generic grasping behavior (Voudouris et al., 2019). If the mass distribution remains invariant over repeated trials, humans rapidly learn and establish more reliable predictions about the object’s properties, which helps to tailor their motor behavior by adopting more suitable grasping postures and by applying more efficient digit forces that foster a stable grasp and object manipulation (Fu et al., 2010). In such cases, it is evident that humans learn the object dynamics through prior experiences that arise from somatosensory feedback during object interactions.

Despite the importance of somatosensory feedback for motor control and learning, somatosensory afferents, in particular those from the tactile domain, are typically suppressed during movement (Beyvers et al., 2022a; Buckingham et al., 2010; Colino & Binsted, 2016; Fraser & Fiehler, 2018; Voudouris et al., 2017; Juravle et al., 2017). This phenomenon of tactile suppression is explained by an internal feed-forward model that predicts future sensory states of the moving limb and suppresses associated sensory signals (Fuehrer et al., 2022; Voss et al., 2008) based on efferent signals related to the underlying movement (Voss et al., 2006; Haggard & Whitford, 2014; Arikan et al., 2021). Tactile suppression is typically assessed by presenting tactile stimuli on a limb when it is moving compared to resting. It has therefore been suggested that peripheral mechanisms may also play some role in tactile suppression (Williams & Chapman, 2002), as proprioceptive afferents arising from the movement itself may mask the perception of the tactile stimulus. However, tactile suppression does not appear to be stronger at moments when sensory feedback signals are stronger (e.g., Broda et al., 2020). In addition, tactile suppression is temporally tuned in line with predictions of an optimal feedback controller (Voudouris & Fiehler, 2021), and is stronger when interacting with predictable than unpredictable environments (Voudouris et al., 2019), supporting the notion that suppression reflects a process related to sensorimotor predictions (Fuehrer et al., 2022).

Sensorimotor predictions are based on a combination of current sensory feedback and prior knowledge about the prevailing dynamics (Wolpert & Flanagan, 2001). When statistical regularities are highly systematic, such as when repeatedly grasping an object of constant mass distribution, sensorimotor predictions are used both to adopt a suitable grasping posture and to suppress associated tactile feedback (Voudouris et al., 2019). Likewise, when the object’s mass distribution changes on a trial-by-trial basis in an unpredictable manner, the adopted grasping configuration remains similar and the associated tactile suppression is substantially weaker (Voudouris et al., 2019). However, even in such unpredictable environments, humans plan their movements on the basis of predictions inferred through their most recent sensorimotor memories (Lukos et al., 2013; Witney et al., 2001). Specifically, people grasp an object of unknown mass distribution *as if it had* the mass distribution experienced in the previous trial. Thus, predictive control seems to be preserved even when acting within unpredictable worlds, and even if these predictions may eventually be incorrect.

In the present study, we sought to examine whether such predictions based on sensorimotor memories influence grasping kinematics and associated tactile processing. We asked participants to grasp and lift an object with a mass distribution that could not be inferred from visual information. The object’s mass distribution changed in a pseudo-randomized order across trials but, unbeknownst to the participants, consecutive trials could involve either the same or different mass distributions. At the moment of object contact, a brief vibrotactile probe stimulus on the index finger of the grasping hand was presented to probe tactile suppression (i.e., Colino & Binsted, 2016; Voudouris et al., 2019) and participants had to report whether they detected that probe stimulus or not. Based on previous work (Lukos et al., 2013), we expected previous trial effects on various kinematic variables. We also expected tactile suppression during grasping compared to rest (Colino & Binsted, 2016; Voudouris et al., 2019). If tactile suppression is modulated by predictions established on the basis of sensorimotor memories, we expected stronger tactile suppression when the mass distribution of an object is repeated from one trial to another compared to when it changes. Alternatively, if tactile suppression is affected by peripheral mechanisms that mask the tactile probe (e.g., Williams & Chapman, 2002), we expect stronger tactile suppression when the actual mass distribution is different from the predicted, because in this case we expect increased processing of afferent information (i.e. Franklin et al., 2012).

## Experiment 1

### Methods

#### Participants

Twenty-four participants (18 female, 6 male) aged 19-34 years (*mean* = 23 ± 3.6) participated in this experiment. The sample size was based on previous studies on tactile suppression (Beyvers et al., 2022a; Fraser & Fiehler, 2018; Fuehrer et al., 2022; Gertz et al., 2018; Manzone et al., 2018; Voudouris et al., 2019). Participants were all right-handed as measured by the German translation of the Edinburgh Handedness Inventory (Oldfield, 1971; *range* = 600-100), free from any known neurological conditions, and had normal or corrected-to-normal visual acuity. All participants gave informed written consent to participate in the experiment and received 8 €/hour or corresponding course credits as compensation for their effort. The experiment was approved by the research ethics board of the Department of Psychology and Sports Science, Justus Liebig University Giessen and was in accordance with the Declaration of Helsinki (2013, except pre-registration of the study).

#### Apparatus

Participants had to grasp and lift an inverted T-shaped object (Figure 1a). The object had a depth of 5 cm. The lower part was 15 cm wide and 3 cm high, while the upper part was 5 cm wide and 7 cm high. At the backside of the lower part of the object, invisible to the participants, three tubes were distributed laterally. A cylindrical piece of brass (116 g) was inserted in either the left or the right tube, creating an asymmetric object’s mass distribution (MD). The total mass of the object, including the brass, was 273 g.

**Figure 1.**
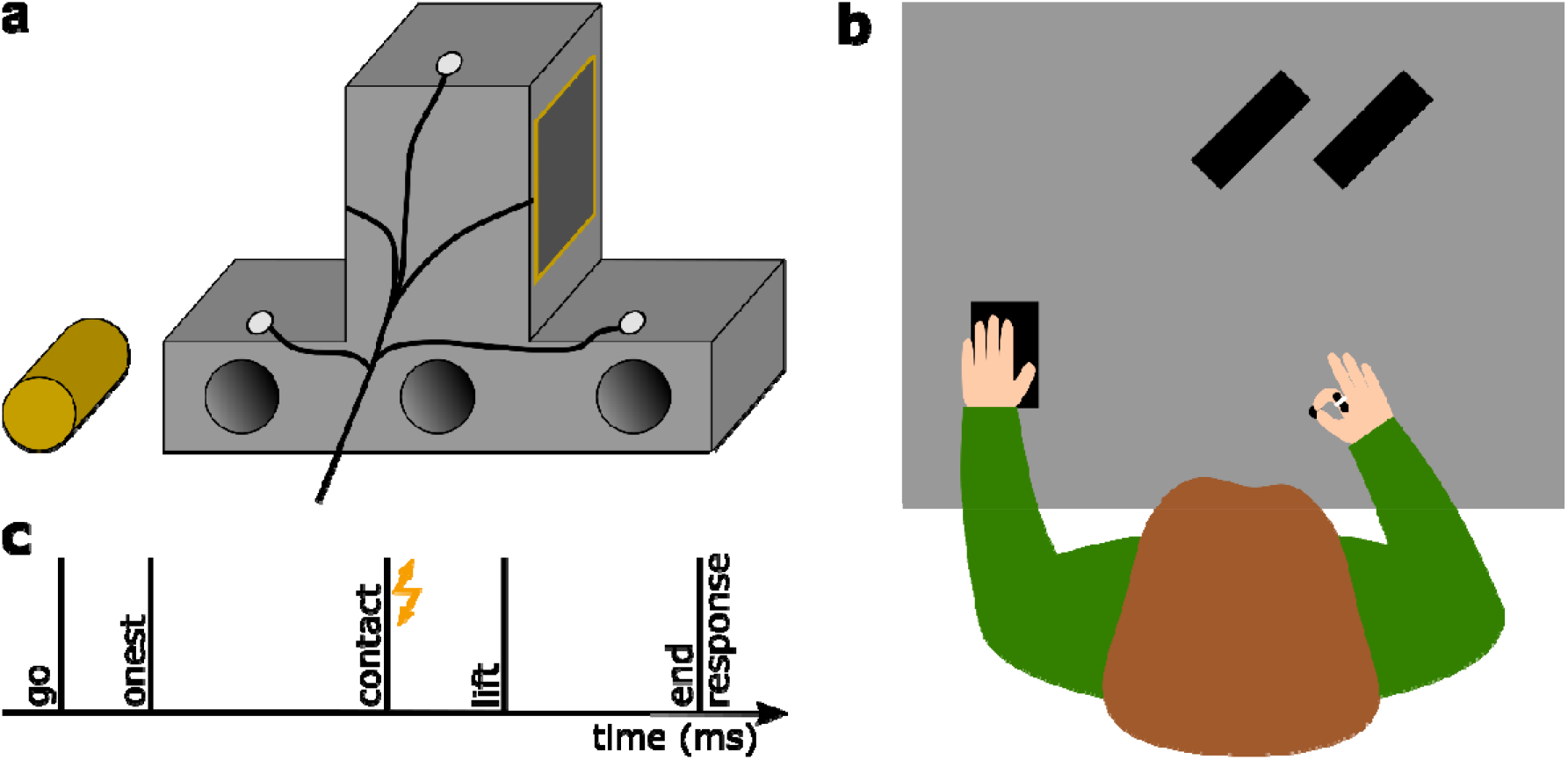
Experimental setup. (a) Illustration of the backside of the inverted T-shaped object with three infrared markers (white circles) and touch sensors (here only one visible) fixed to the grasping area of the object. The three tubes in the object’s base were never visible to the participant. The brass mass is depicted to the left of the object. (b) Top view of the setup with the right index finger and thumb resting on the start position and the left hand resting on the numpad. A vibrotactile sensor was fixed on the dorsal part of the participant’s right index finger. Both object positions are also depicted (black rectangles). (c) Timeline of a single trial with the flash illustrating the vibrotactile stimulation at the moment of object contact.

Participants were seated comfortably in front of a 117 cm × 80 cm table (Figure 1b). A small keypad (12.5 cm × 8 cm) on the table was under the participant’s left hand at a comfortable position. The start position for each reaching-to-grasp movement was aligned to the participants’ right shoulder and was marked by a round piece of felt pad located ∼30cm in front of their body. To avoid stereotyped movements, two different lateral object positions were used, each 34 cm from the start position. To prevent participants from seeing in which tube the experimenter inserted the brass prior to each trial, participants wore liquid-crystal shutter glasses (PLATO, Translucent Technologies, Toronto, Canada) throughout the grasping block (see below).

To track the movement of the participants’ hand and of the object, the position of five infrared markers were recorded at 100 Hz with an Optotrak Certus motion tracking system (Northern Digital Inc., Waterloo, ON, Canada). One marker each was attached to the fingernail of the participants’ right thumb and index finger. The other three markers were attached in a triangular arrangement on the backside of the object.

Vibrotactile stimulations, always given at the moment of object touch, were triggered by a touch sensor (4.37cm × 4.37 cm; Interlink Electronics Inc., Westlake Village, CA, USA) that was mounted on each grasping side of the upper part of the object and that was connected through a NI USB-6009 device (National Instruments Corporation, Austin, TX, USA) to the host PC. A vibrotactile stimulation device (Engineering Acoustics Inc., Casselberry, FL, USA) was attached to the dorsal part of the medial phalanx of the participants’ right index finger. The presented vibrotactile stimuli (250 Hz, 50 ms) differed in intensity; six stimuli had a peak-to-peak displacement ranging from 9.4 to 56.7 µm in steps of 9.4 µm, while one third of the trials involved no stimulation. Care was taken to ensure that motion tracking markers, the tactor and the connected cables did not constrain the participants’ freedom of movement.

#### Procedure

The experiment consisted of three blocks: two baseline blocks and one grasping block. In the baseline blocks, participants rested their right hand in a comfortable position in front of them. Each baseline trial started automatically: A probe vibrotactile stimulus was presented, if at all, 500 ms after the beginning of the trial. An auditory cue 500 ms after that moment prompted participants to press a button underneath their left index finger or thumb and indicate whether they felt a vibration or not, respectively. The next trial started automatically 500 ms after the response was given. During each of the two baseline blocks, each of the six stimulus intensities was presented five times, while the no-stimulation catch trials were presented 15 times, resulting in a total of 45 trials presented in a pseudorandom order. One baseline block each was presented before and after the grasping block in order to account for possible changes in tactile sensitivity over time.

During the grasping block, each trial started with the shutter glasses being opaque and the participant bringing their right thumb and index fingertips together at the start position. The experimenter inserted the brass mass into the appropriate tube and then placed the object on the appropriate position. Even if the mass distribution was similar between two consecutive trials, the brass was always pulled out and inserted into the given tube to keep auditory cues constant. After the experimenter pressed a key to start the trial, the shutter glasses turned transparent and an auditory signal indicated that participants could start their movement. They were to reach out and grasp the object with their right thumb and index finger at both sides of its upper part, where the touch sensors were attached, lift it as straight as possible for ∼10 cm, before placing it back down on the table and returning their hand to the start position. Once participants first touched the object, as this was detected by the first activation of the touch sensors, a probe vibrotactile stimulus could be presented on the participant’s right index finger. These probe stimuli were identical to those presented in the baseline blocks. After five seconds following the auditory signal, data collection was stopped, the shutter glasses turned opaque, and an auditory cue prompted participants to indicate whether they had felt a probe stimulus (yes/no response). Participants responded in the same way as in the previously described baseline procedure. Figure 1c illustrates the progression of a single grasping trial.

In the grasping block, each participant was presented with a trial sequence of mixed mass distributions. This was designed in a way that all possible combinations of consecutive mass distributions occurred equally often during the block: left followed by left (LL), right followed by left (RL), left followed by right (LR), and right followed by right (RR). For each of these four combinations, 30 trials with vibrotactile stimuli and 15 without a stimulation were presented, identical to those presented in one baseline block. This mixed sequence consisted of trials where the mass distribution repeated (*mixed*_*same*_; LL and RR) or changed (*mixed*_*different*_; LR and RL) from one trial to the next. Overall a total of 180 trials were presented, with 90 trials having the object on the left and the other 90 trials on the right target position in a pseudorandom order across trials. Before starting the grasping block, participants performed nine practice trials to familiarize with the task and the object weight. For those trials, the brass cylinder was placed into the central tube of the object, and this was the only case when the symmetric mass distribution was used. In these practice trials, each stimulus intensity was presented once, in addition to three catch trials with no stimulation. The object was randomly placed five times on the left position and four times on the right position, or the other way around, counterbalanced across participant.

#### Data analysis

The first goal of the study was to examine whether participants plan their upcoming movement on the basis of sensorimotor memories, i.e. sensorimotor information experienced in the previous trial. To analyze the relevant kinematic behavior, we examined the influence of the previous trial’s MD on (a) the vertical separation of the digits at the moment of first object contact, (b) the time between first object contact and object lift, and (c) the maximal object roll during lifting. To determine the relevant kinematic measures for each trial, we first calculated the speed of the hand and of the object by numerical differentiation of the mean position of the markers on both digits and on the object, respectively. These two speed vectors were dual-pass filtered using the MATLAB *filtfilt* function with a 2nd order lowpass butterworth filter and a cutoff frequency of 0.3 Hz. Object lift onset was defined as the first timepoint at which the object speed was greater than 10 cm/s. The moment of object contact was determined as the last timepoint before object lift onset with a hand speed lower than 10 cm/s. For each trial, we calculated the *digits’ separation* as the mean vertical distance between the thumb and the index finger within the first 100 ms from the moment of object contact, with positive values indicating that the thumb was placed higher than the index finger (Voudouris et al., 2019). *Loading time* was defined as the time between the moment of contact and object lift onset. Finally, *object roll* was defined as the absolute maximal tilt angle within the first 250 ms after the onset of the object lift.

Next, we examined whether kinematic behavior on a trial with a given MD was influenced by the MD experienced in the previous trial. We defined two configurations of consecutive MDs: *MD*_*same*_ involved consecutive MDs that were identical (LL, RR), whereas *MD*_*different*_ involved different consecutive MDs (RL, LR). For *digits’ separation*, we first calculated four average values per participant, one across the trials of each of the four possible combinations. To avoid negating the actual effect, the average digits’ separation in trials with left MD were subtracted from the corresponding value in trials with right MD, separately for *MD*_*same*_ (RR – LL) and *MD*_*different*_ (LR – RL) configurations. Based on previous work (Lukos et al., 2013; Voudouris et al., 2019), we expected participants to place their thumb lower and higher than the index finger when they anticipated the mass to be on the right and left side, respectively, which would minimize object roll during object lift. As these would lead to negative and positive digits’ separation indices (see above), the *MD*_*same*_ calculation (RR – LL) should yield negative indices and the *MD*_*different*_ calculation (LR – RL) should yield positive indices. In other words, if the resulting index is negative when the mass distribution is repeated and positive when the mass distribution is switched, this should indicate that participants use predictive control and adopt in trial *n* a grasp configuration to match the dynamics experienced in trial *n-1*, independently of whether the adopted grasping posture is suitable for the mass distribution of the grasped object. *Loading time and maximal object roll* were calculated based on the average values that were obtained for each trial for both configurations (*MD*_*same*,_ *MD*_*different*_).

The second goal of the study was to examine whether sensorimotor memories influence somatosensory processing on the grasping hand. In accordance with previous work (Colino & Binsted, 2016; Voudouris et al., 2019), we expected decreased sensitivity to the probing vibrotactile stimuli during grasping compared to the resting trials. If participants consider the MD experienced in the previous trial in order to predict the MD in the upcoming trial, tactile sensitivity should change accordingly: Tactile suppression should be stronger in the second of two consecutive trials with identical MD, as the predicted and experienced movement dynamics match. Alternatively, if tactile suppression is affected by backward masking, we expect stronger suppression in trials with different MD than predicted, due to increased feedback processing (i.e. Franklin et al., 2012). To assess tactile suppression, we first merged the responses of both baseline blocks and fitted each participant’s responses in each condition (baseline, *MD*_*same*_, *MD*_*different*_) with separate logistic functions using maximum-likelihood estimation with the function *psignifit 3* (Wichmann & Hill, 2001) in MATLAB 2019b (MathWorks, Natick, MA). For each of these three psychometric functions that we obtained for each participant, we estimated the detection threshold as the stimulus intensity at 50% of the function. To quantify tactile suppression during grasping while accounting for individual tactile sensitivity, each participant’s baseline detection threshold was subtracted from their respective values of the two grasping configurations (*MD*_*same*_, *MD*_*different*_). Each of these suppression values were calculated for each participant, and then averaged across participants, with higher positive values indicating stronger tactile suppression. Participants only took part in the whole experiment if their false positive rate in the baseline trials was below 30%. In the grasping block, false alarm rates were below 30% for all participants. Further, we merged the two baseline blocks because we found no differences in the respective detection thresholds.

After calculating the above-mentioned variables, we examined effects of sensorimotor memories on grasping kinematics and associated tactile suppression. For kinematic behavior, we submitted the index values for digits’ separation to two-sided one sample t-tests against zero, which tested whether participants adopted a grasping posture based on the mass distribution experienced in the trial before. For tactile processing, we examined whether detection thresholds during each grasping condition was greater than those during baseline by submitting the suppression scores in two-sided one sample t-tests against zero. In a second level, we investigated the effect of previous mass distribution on all kinematic variables and tactile suppression, by using separate paired samples t-tests for grasping kinematics and tactile processing between trials with repeating (*MD*_*same*_) and changing MD (*MD*_*different*_). Effect sizes are described as Cohen’s d. All statistical analyses were carried out with JASP (Version 0.14.1).

## Results

### Kinematics

Regarding finger positioning on the object, we found mean index values below zero for the *MD*_*same*_ condition (*t*_*23*_ = −5.07, *p* < .001, *d* = −1.04) and above zero for the *MD*_*different*_ condition (*t*_*23*_ = 8.08, *p* < .001, *d* = 1.65), indicating that participants positioned their fingers in anticipation of a MD that was identical to the one presented in the trial before, independently of whether this prediction was correct or not. Accordingly, the index value was smaller for *MD*_*same*_ than *MD*_*different*_ (*t*_*23*_ = −10.83, *p* < .001, *d* = −2.21; Figure 2a).

**Figure 2.**
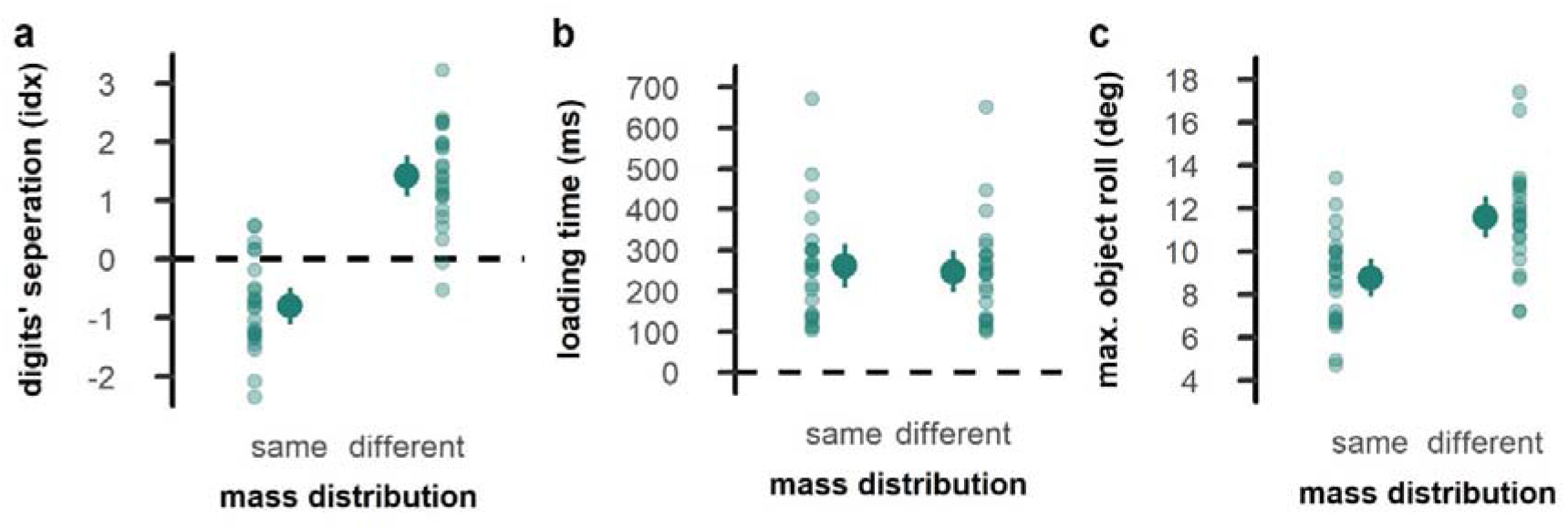
Kinematic performance in Experiment 1. (a) Digits’ vertical separation at the moment of object contact, (b) loading time, and (c) maximal object roll during the first 250 ms after object lift. The left side of each panel shows trials with a repeated mass distribution, and the right side for trials with a changing mass distribution. Opaque dots represent the mean across participants with error bars indicating the confidence interval. Transparent dots represent individual participants.

The MD experienced in the previous trial also affected loading time (*t*_*23*_ = 5.42, *p* < .001, *d* = 1.11; Figure 2b) and maximal object roll (*t*_*23*_ = −18.93, *p* < .001, *d* = −3.87; Figure 2c) in the current trial. Specifically, loading times were longer and object roll was smaller when the MD was repeated from one trial to another compared to when it was changed.

### Tactile sensitivity

As expected (Colino & Binsted, 2016; Voudouris et al., 2019), tactile suppression was evident during grasping compared to rest. This was the case for both the *MD*_*same*_ (*t*_*23*_ = 7.04, *p* < .001, *d* = 1.44) and *MD*_*different*_ (*t*_*23*_ = 8.452, *p* < .001, *d* = 1.72). Yet, we found no effects of previous trial MD on tactile suppression (*t*_*23*_ = .33, *p* = .742, *d* = .07; Figure 3).

**Figure 3.**
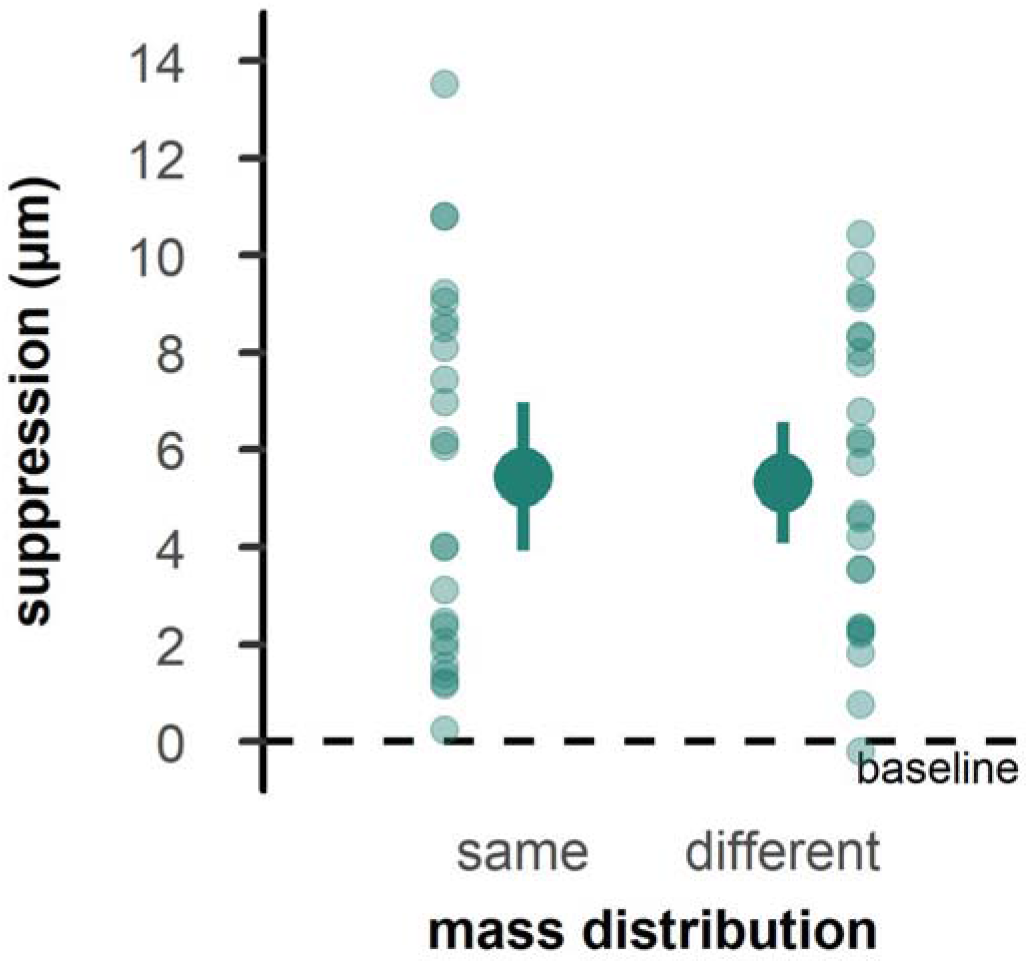
Tactile suppression scores in Experiment 1. The left side of each panel shows trials with a repeated mass distribution, and the right side trials with a changing mass distribution. Zero values indicate no change from baseline, while higher values indicate a deterioration of tactile perception. Details as in Figure 2.

## Discussion Experiment 1

We investigated whether and how sensorimotor memories influence grasping kinematics and associated tactile processing. Our results show that, when grasping an object with uncertain properties, humans utilize sensorimotor memories based on previous trial experience to plan their next movement (cf., Lukos et al., 2013). Specifically, participants grasped an object of unknown mass distribution by predictively choosing contact points that would compensate the mass distribution experienced in the *previous* trial, even if the mass distribution in the current trial was different than predicted. These results suggest that sensorimotor memory is used to adjust movement kinematics when object properties are uncertain and sensory action consequences are hard to predict. In line with previous findings (Colino & Binsted, 2016; Voudouris et al., 2019), tactile sensitivity was decreased during grasping compared rest reflecting movement-induced tactile suppression. However, tactile suppression was unaffected by the mass distribution of the previous trial, as it did not differ between consecutive trials of repeating and changing mass distributions. These results suggest that tactile suppression is neither affected by predictions based on sensorimotor memories (*MD*_*same*_ > *MD*_*different*_), nor by backward masking (*MD*_*same*_ < *MD*_*different*_).

In this experiment, participants had no information about the mass distribution as this could change from trial to trial. We also probed tactile suppression at a moment when the current mass distribution could not be confirmed. Thus, any predictions based on sensorimotor memories might have been equally strong for trials that involved the same (*MD*_*same*_) compared to trials that involved a different mass distribution than the one predicted (*MD*_*different*_). To address this possibility, we conducted Experiment 2, where we interleaved longer sequences of trials involving an object of constant mass distribution with shorter sequences of trials with changing mass distribution. We assumed stronger sensorimotor predictions in sequences of constant compared to mixed distributions, and thus expected stronger suppression in those sequences. This experiment was further split into two sessions: in the first session, participants were exposed to the interleaved constant and mixed sequences to test for implicit learning of the statistical regularities of the trial sequence. In the second session, the experimenter informed the participants explicitly about the upcoming sequence (constant or mixed). This was done to explore whether explicit information about the upcoming trial sequence would alter predictive control and the strength of tactile suppression with stronger suppression for the constant compared to the mixed sequence.

## Experiment 2

### Methods

#### Participants and Apparatus

Twenty-four participants (14 female, 10 male) aged 18-31 years (*mean* = 22.71 ± 3.46) participated in the study and completed the experiment. None of the participants took part in Experiment 1. They were all right-handed as measured by the Edinburgh handedness inventory (range = 60-100) and had normal or corrected-to-normal visual acuity. As in Experiment 1, participants gave their written informed consent and received monetary compensation or course-credits for their efforts. The setup, procedure and analyses were identical to those of Experiment 1, except for the details mentioned below.

#### Procedure

Experiment 2 consisted of two sessions, each performed on different days. Each session included two baseline blocks, a short practice and a grasping block, all following the same procedure as in Experiment 1. The main difference was the implemented trial sequence. We created a trial sequence with longer parts of 27 trials with a constant MD (*constant*), which were interrupted by short parts of nine trials with a pseudorandomized MD (*mixed*). The longer constant parts should allow building more stable predictions. The mixed interruptions were kept short to limit the total duration of the experiment allowing it to be completed in one session. Since we did not find an effect of the previous trial’s MD on tactile suppression in Experiment 1, we did not split the mixed sequence in *MD*_*same*_ and *MD*_*different*_ trials. Rather, we compared suppression only between mixed and constant parts. We presented three constant parts with a left MD and three blocks with a right MD, always interrupted by a mixed part. The sequence always started with a mixed part and consisted of 216 trials in total. The mixed parts were constructed in the same way as the mixed sequence in Experiment 1. Within the constant, longer parts, one of the two possible MDs was chosen, counterbalanced across participants, but the object position could still change within these parts. During the first session, participants were not informed about the structure of the presented sequence (*implicit version*). At the end of this session, participants filled out a custom questionnaire asking whether they observed a specific trial structure of the presented sequence. Only one of the twenty-four participants indicated that they noted some repetitions. During the second session, performed on average seventeen days later, the very same sequence was presented to each participant. During this session, the experimenter explicitly informed the participant verbally about the structure of the upcoming part of the sequence by indicating whether they would encounter a mixed or constant part (*explicit version*), without however indicating what mass distribution would be presented in the upcoming constant part.

#### Data Analysis

Conditions were divided according to the parts of the sequence (mixed, constant) and the experimental version (implicit, explicit). The effects of sequence part (mixed, constant) and experimental version (implicit, explicit) on digits’ separation, loading time, object roll, and tactile suppression were evaluated using separate 2 × 2 repeated measures ANOVAs. Significant interactions were inspected with post-hoc t-tests that were Bonferroni-corrected for multiple comparisons. Effect sizes are described as partial Eta squared 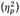 for ANOVAs and as Cohen’s d for t-tests.

## Results

### Kinematics

For digits’ separation, a negative index occurring when the mass distribution is repeated and a positive when the mass distribution is switched, indicate that participants adopt in trial *n* a grasp configuration to match the dynamics experienced in trial *n-1*, independently of whether the adopted grasping posture is suitable for the mass distribution of the grasped object. As expected, and in line with the results of Experiment 1, in the constant parts participants positioned their fingers in anticipation of a repeated MD in both the implicit (*t*_*23*_ = −6.23, *p* < .001, *d* = −1.27) and the explicit version of the experiment (*t*_*23*_ = −6.53, *p* < .001, *d* = −1.33). Likewise, in the mixed parts participants positioned their fingers in anticipation of a repeated MD, irrespective of whether the MD was repeated or changed. This was the case during both the implicit (*t*_*23*_ = 2.39, *p* =.026, *d* = .49) and the explicit version (*t*_*23*_ = 3.51, *p* = .002, *d* = .72). Evidently, these resulted in a main effect of sequence part (*F*_1, 23_ = 61.12, *p* < .001,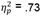; Figure 4a). We found no differences between implicit and explicit parts, nor an interaction (both *F* < 1.3, both *p* > .30, both 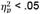).

**Figure 4.**
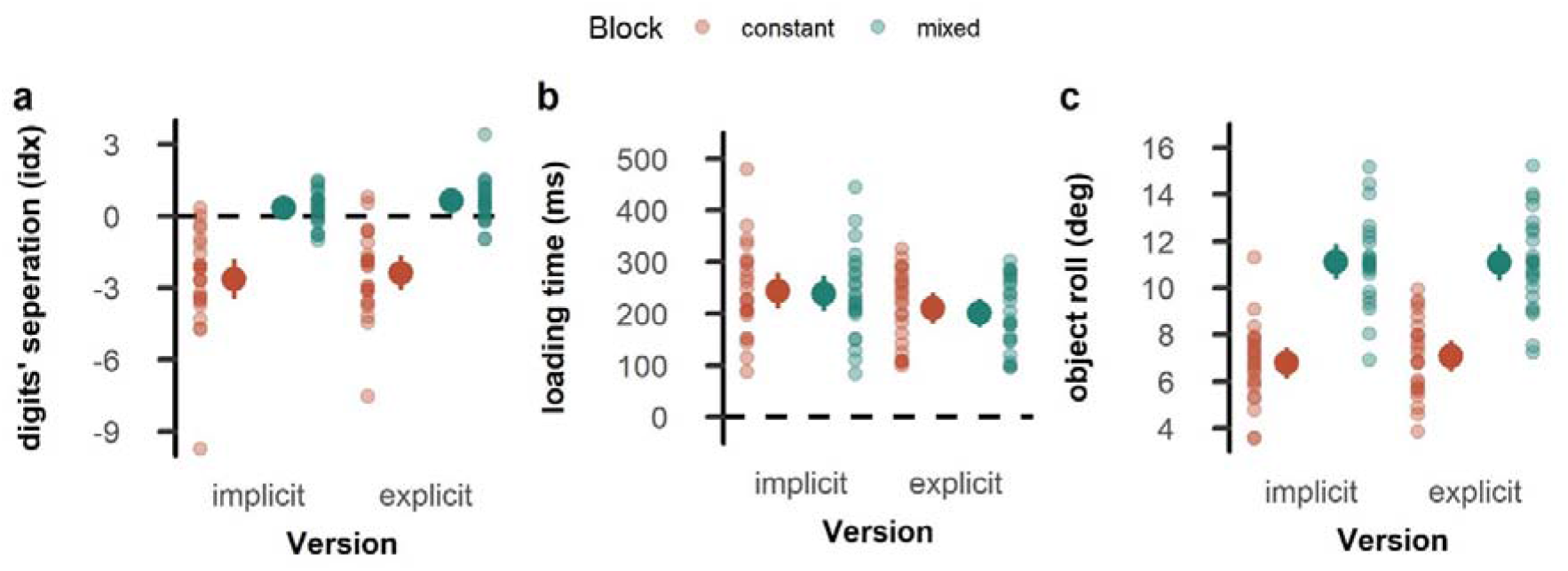
Kinematic performance in Experiment 2. (a) Digits’ vertical separation at the moment of object contact, (b) loading time, and (c) maximal object roll during the first 250 ms after object lift. The left side of each panel shows results for the implicit version, and the right side for the explicit version. Mean values for the constant blocks are depicted in red, those for the mixed blocks in green. Opaque dots represent the mean across participants with error bars indicating the confidence interval. Transparent dots represent individual participants.

Participants took longer to start lifting the object when the MD was constant over several trials compared to when the MD changed (*F*_*1, 23*_ = 6.82, *p* = .016, 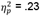). They also took longer to start lifting the object in the first, implicit version of the experiment, compared to the second, explicit version (*F*_*1, 23*_ = 11.56, *p* = .002,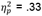; Figure 4b). There was no interaction between the two factors (*F*_1, 23_ = .70, *p* = .411,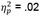). Object roll was smaller during the lift when the MD was repeated over several trials compared to when the MD changed (*F*_1, 23_ = 470.69, *p* < .001,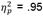; Figure 4c), in line with the results of Experiment 1. There was no effect of the experimental version nor an interaction (both *F* < 2.5, both *p* > .13, both 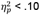).

### Tactile sensitivity

Tactile suppression was evident during grasping compared to rest in the implicit version both in the constant (*t*_*23*_ = 7.15, *p* < .001, *d* = 1.46) and the mixed parts of the sequence (*t*_*23*_ = 8.72, *p* < .001, *d* = 1.78), as well as in the explicit version, again both in the constant (*t*_*23*_ = 7.41, *p* < .001, *d* = 1.51) and the mixed part of the sequence (*t*_*23*_ = 7.47, *p* < .001, *d* = 1.52). Suppression appeared stronger in the mixed than constant parts, but this difference was not statistically significant (*F*_*1, 23*_ = 4.01, *p* = .057,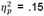). There was also no difference in suppression between implicit and explicit versions (*F*_*1, 23*_ = 0.45, *p* = .510, 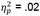; Figure 5), indicating that the type of knowledge about the statistical regularities of the upcoming trial sequence did not systematically affect tactile suppression. There was also no interaction between the factors (*F*_*1, 23*_ = 2.69, *p* = 0.114, 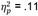).

**Figure 5.**
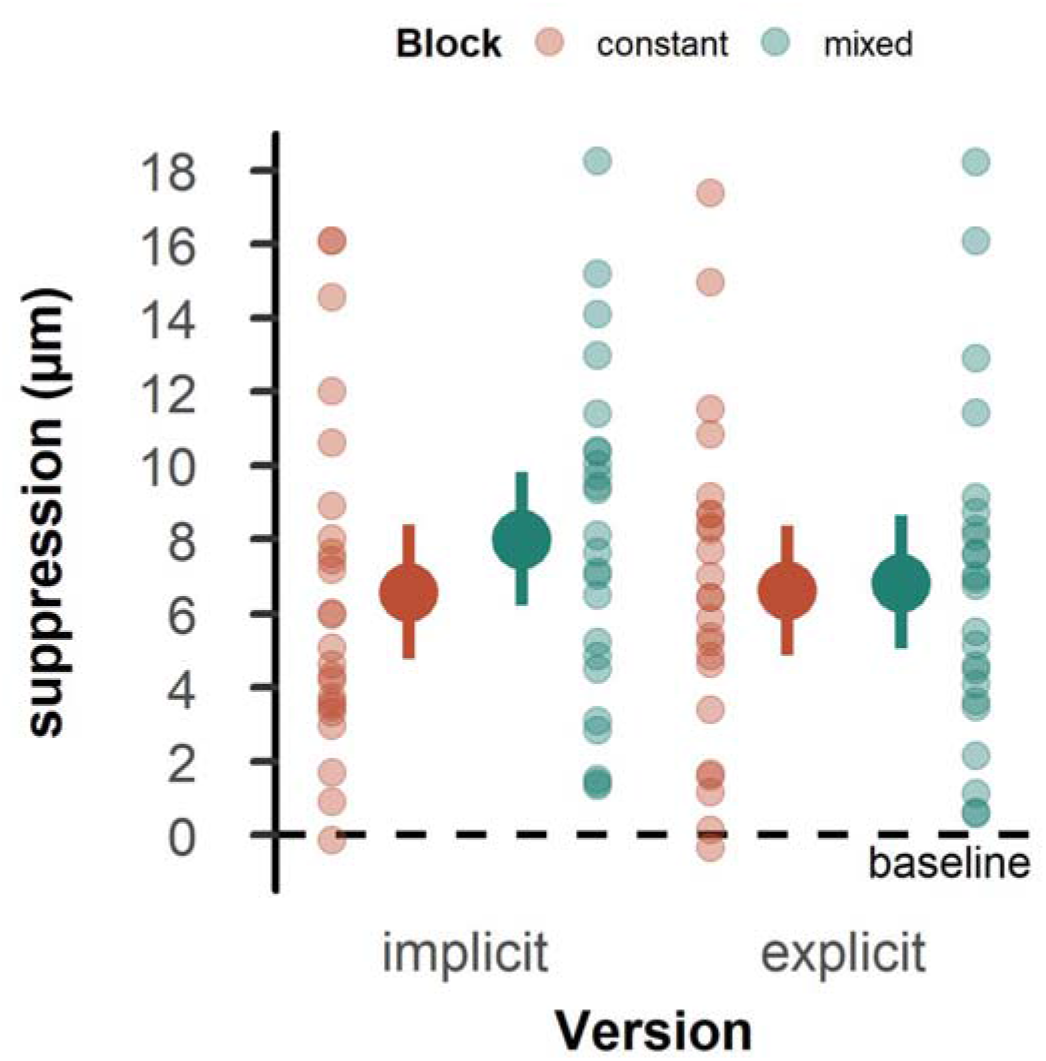
Tactile suppression scores in Experiment 2. The left side of each panel shows the implicit version, and the right side shows the explicit version. Mean values for the constant blocks are depicted in red, those for the mixed blocks in green. Zero values indicate no change from baseline, while higher values indicate a deterioration of tactile perception. Opaque dots represent the mean across participants with error bars indicating the confidence interval. Transparent dots represent individual participants.

## Discussion Experiment 2

In Experiment 2 we investigated whether longer parts of repeated object properties would lead to more reliable sensorimotor predictions and hence stronger tactile suppression. Further, we were interested in whether implicit or explicit information about the trial structure would affect tactile suppression. Grasping kinematic were affected similarly to Experiment 1, with grasping postures being predictively chosen to compensate for object roll, even if this prediction might have been wrong. We again found tactile suppression during grasping compared to rest, but suppression was similar between the constant and mixed parts. These results support and further extend the findings of Experiment 1, showing that sensorimotor memories influence movement kinematics but not associated tactile perception. The similar suppression between constant and mixed parts is, however, at odds with previous findings, where suppression was stronger when grasping an object whose mass distribution remained identical over trials than when it changed (Voudouris et al., 2019). One possible explanation is that in the current study, participants were exposed to the constant parts for shorter durations and these constant parts were alternating with mixed parts, which might have limited the establishment of reliable predictions that could affect tactile suppression.

Implicit and explicit information about the upcoming trial structure did not affect the grasping posture or object roll, and it had no effect on tactile suppression. Yet, our results showed that during the explicit session of the experiment, participants started lifting the object earlier than during the implicit session. We interpret this finding with caution, as the explicit version was presented always *after* the implicit version. This does not allow us to conclude whether this effect is caused by the explicit information itself or whether it is a byproduct of practice, as participants may have lifted the object faster simply because they were more familiar with the task.

## General Discussion

In this study we examined whether sensorimotor memories influence grasping kinematics and associated tactile processing. Participants reached to grasp an object that could have one of two possible mass distributions. When the mass distributions were presented in a pseudorandom order across trials (Experiment 1), participants grasped the object in the second of two consecutive trials as if it had the mass distribution experienced in the previous trial, demonstrating predictive grasping behavior based on sensorimotor memories. Similar behavior was observed in Experiment 2, when the mass distributions were presented both in a mixed and in a constant order. To assess tactile processing, we employed the well-established phenomenon of tactile suppression and found clear suppression of tactile signals during grasping compared to rest. However, the strength of tactile suppression was unaffected by the statistical regularities of the trial sequence. Our results indicate that sensorimotor memories exert a strong influence on movement behavior but not on associated tactile processing.

Goal-directed behavior is primarily based on visual information. For instance, when grasping an object, humans use vision to extract relevant object properties, such as its size (Gordon et al., 1991), shape (Kleinholdermann et al., 2007), orientation (Voudouris et al., 2013), or surface material (Klein et al., 2021), and they choose suitable grasping points already during movement planning (Roche et al., 2015; Voudouris et al., 2010). However, visual information is not always available or informative about certain object properties. For instance, the object’s center of mass may not always be inferred from vision. When interacting with such objects, humans initially adopt a ‘default’ motor behavior, and if object properties remain invariant and occur recurrently, they use somatosensory feedback from the previous interaction to predict the object properties and tailor their movement accordingly (Fu et al., 2010). However, when object properties change continuously, predictions are less reliable and can be misleading. Yet, even in such cases, humans plan their movements assuming that the object in question has the same properties as those experienced in the previous trial (Beyvers et al., 2022b; Lukos et al., 2013; Witney et al., 2001). Our experiments support such predictive behavior. Specifically, participants grasped the object in the second of two consecutive trials as if it had the dynamics experienced in the trial before. Here, the posture cannot have been chosen based on somatosensory feedback from the object of the current trial because grasping posture is assessed before such feedback could be utilized. This tuning of grasping posture demonstrates that motor plans are established based on sensorimotor memories. Evidently, such memories can be advantageous when the prediction is correct but disadvantageous when it is incorrect, as reflected in object roll during object lift.

The second aim of the study was to examine whether tactile processing associated with object grasping and manipulation is affected by predictions established through sensorimotor memories. The results of our two experiments demonstrate robust tactile suppression during movement, in line with previous findings (Colino & Binsted, 2016; Voudouris et al., 2019). This supports the idea that sensorimotor predictions established during movement reduce associated tactile sensitivity (e.g., Fuehrer et al., 2022). In Experiment 1 we show that tactile suppression did not differ between trials of the same than different mass distribution compared to the distribution experienced in the previous trial. This suggests that, although predictions from sensorimotor memories influence grasping kinematics, these predictions do not affect tactile suppression. One possible explanation for this finding is that we assessed tactile suppression at the moment of first contact with the object. At that moment, participants had no access to sensory input about the actual mass distribution, and the sensorimotor prediction might have been equally strong, independently of the object’s actual mass distribution. In such case, it is unsurprising that tactile suppression is similar between conditions. In Experiment 2 we included a condition with sequences of 27 consecutive trials that involved an object of constant mass distribution. Such constant parts should facilitate predictions about the object’s mass distribution compared to the mixed sequence, in which the mass distribution was changing on a trial-by-trial basis. Despite the long sequences of trial repetitions in the constant parts, tactile suppression was similar between constant and mixed parts. This is at odds with previous findings that demonstrated stronger suppression in constant than mixed sequence parts (Voudouris et al., 2019). Such difference may arise due to the fact that in our current experiment the mixed and constant parts were relatively short and were presented interchangeably, whereas in the other study the mixed and constant parts were much longer and were presented in distinct blocks. Thus, in the present study, tactile suppression might not have been adapted as effectively, as participants had to constantly adjust to a switch between constant and mixed intercepts, and as a consequence the sensorimotor predictions in the constant parts might have been weaker compared to the previous study (Voudouris et al., 2019).

Tactile suppression has also been explained by peripheral mechanisms, such as backward masking from movement-related afferent signals that mask the tactile probe used to measure suppression (e.g., Williams & Chapman, 2002). In this case, we would have expected stronger suppression in trials where the prediction about the object’s mass distribution was wrong, such as in *MD*_*different*_ than *MD*_*same*_ trials of Experiment 1, because when experiencing an unpredictable sensory event during goal-directed actions, feedback gains increase (e.g., Franklin et al., 2012). In Experiment 2, it appears that suppression is qualitatively stronger in the mixed compared to the constant condition of the implicit version, however this was not systematic. All in all, our results do not lend support for the hypothesis of backward masking either, and the apparent invariance in suppression as a function of feedback signals is in line with previous findings (Broda et al., 2020).

In Experiment 2 we further examined whether grasping kinematics and tactile suppression are sensitive to statistical regularities that are implicitly or explicitly inferred. Previous research has demonstrated that implicit learning of the underlying dynamics can be beneficial for sampling, for instance through haptic exploration (Zoeller et al., 2019). In addition, implicit sensorimotor memories of object properties can influence the anticipation of these properties (Schneider et al., 2020), both in the absence and presence of explicit cues (Flanagan et al., 2008). Yet our kinematic results did not reveal any benefit from the explicit instruction about the trial sequence. Albeit, we observed a slight decrease of loading times when explicit information about the upcoming sequence was provided. However, it is noteworthy that this benefit may simply be due to the repetitive performance of the task, as the explicit session was *always* presented after the implicit session, so we cannot disentangle effects of practice from effects of the type of information. Importantly, we also did not observe any effects of implicit or explicit information on tactile suppression. Informing participants about the fact that the upcoming trials would be of a mixed or a constant sequence does not have any strong effect on motor behavior or associated tactile processing.

In conclusion, we demonstrate that predictions based on sensorimotor memories influence grasping kinematic by adjusting the grip to the previously experienced object properties. However, associated tactile suppression appears robust to such previous trial effects.

## Acknowledgments

This work was supported by the German Research Foundation (DFG) through the IRTG 1901 “The Brain in Action” as well as through the Collaborative Research Center SFB/TRR 135, project A4, under grant agreement 222641018. It is also supported by “The Adaptive Mind” funded by the Excellence Program of the Hessian Ministry for Higher Education,n Research, Science and the Arts. We would like to thank Friederike Weissmann and Nikola Zalomska for their assistance with participant recruitment and data collection for this work. The data collected for this work will be publicly available at https://doi.org/10.17605/osf.io/w6ruy.

## Competing interests

The authors have no competing interests to declare.

## Authors contributions

M.C.B.: Conceptualization, Methodology, Software, Validation, Formal analysis, Investigation, Data Curation, Writing - Original Draft, Writing - Review & Editing, Visualization, Project administration

D.V.: Conceptualization, Methodology, Writing - Review & Editing, Supervision, Project administration

K.F.: Conceptualization, Methodology, Resources, Writing - Review & Editing, Supervision, Project administration, Funding acquisition

## Notes

### Competing Interest Statement

The authors have declared no competing interest.

